# Selection-driven tumor evolution involving non-cell growth promotion leads to patterns of clonal expansion consistent with neutrality interpretation

**DOI:** 10.1101/2020.02.11.944843

**Authors:** Jack Edwards, Andriy Marusyk, David Basanta

## Abstract

Cancers are the result of eco-evolutionary processes fueled by heritable phenotypic diversification and driven by environmentally dependent selection. As space represents a key growth-limiting ecological resource, the ability to gain and explore this resource is likely to be under strong selection. Using agent-based computational modeling, we explored the consequences of the interplay between phenotypic strategies centered on gaining access to new space through cell-extrinsic degradation of extracellular matrix barriers and the exploitation of this resource through maximizing cell proliferation. While cell proliferation is a cell-intrinsic property, newly accessed space represents a public good, which can benefit both producers and non-producers. We found that that this interplay results in ecological succession, enabling emergence of large, heterogenous, and highly proliferative populations. Even though in our simulations both remodeling and proliferation strategies were under strong positive selection, their interplay led to sub-clonal architecture that could be interpreted as evidence for neutral evolution, warranting cautious interpretation of inferences from sequencing of cancer genomes.

## Introduction

Cancer starts when a somatic cell breaks away from the rules of homeostatic cooperation imposed by multicellularity and starts acting as an independent evolutionary entity. As this cell divides, its progeny (clone) acquires genetic and epigenetic alterations generating novel phenotypic variants (subclones). This subclonal diversification in the context of competition for growth-limiting resources between cancer cells enables Darwinian evolution, which can lead to the emergence of key tumor phenotypes, defined by hallmarks of cancers, culminating in lethal malignancies (Axelrod et al., 2006; Greaves and Maley, 2012; Hanahan and Weinberg, 2011, 2000; Korolev et al., 2014; Nowell, 1976). Consistently, genetic and phenotypic Intratumor Heterogeneity (ITH) correlates with poor prognosis and contributes to treatment failure (Gerlinger et al., 2012; Swanton, 2012). Understanding the basic rules that govern the somatic evolution and how ITH is maintained in the face of subclonal competition is thus key to a more effective tumor prevention and therapies (Basanta and Anderson, 2013; Sottoriva et al., 2015).

Mathematical modelling has been used to understand the behavior of complex dynamic systems, such as changes in cell populations in space and time during the emergence and progression of cancers, which are difficult to intuit without the help from rigorous theoretical frameworks. Properly parameterized and experimentally tested mathematical models can provide a deeper understanding of these complex processes and also serve as predictive tools for therapy optimization. Ordinary Differential Equations (ODE), the most commonly used tools to model somatic evolution, typically assume well-mixed populations (Altrock et al., 2015; Gatenby and Gillies, 2008). This assumption fails to account for heterogeneity in space and tissue architecture as a key growth limiting resource, which has a profound impact on evolutionary dynamics. Agent-based models (ABM), which can consider cells as individual agents rather than populations, can be used to account for space and spatial structures. The importance of space and the applicability of ABMs has been explored showing that an increase in available space results in higher ITH and can affect evolutionary mode (Chkhaidze et al., 2019; Noble et al., 2019; West et al., 2019).

Epithelial tissues, which give rise to the majority of human cancers, are organized as layered structures, constrained by basal membranes and extracellular matrix (ECM) (Bissell et al., 2002; Bissell and Radisky, 2001). Carcinogenesis involves breaking from the growth limitations imposed upon by epithelial layers and gaining access to the new space through the degradation and remodeling of ECM (Cawston and Wilson, 2006). From an ecological perspective, accessing this new space constitutes niche engineering, thus we will refer to the phenotypes of cancer cells which possess the ability to remodel ECM as **engineers** (Lloyd et al., 2016; Myers et al., 2020). ECM degradation is achieved through the secretion of matrix-degrading enzymes, which require a sufficiently high local concentration to achieve their effects. Since the newly accessed space can benefit not only engineers, but also non-engineer cells, the space, unlocked by engineers, represents a *common good* resource (Kagel and Roth, 1995). Optimization for use of space can be achieved by the increased consumption of this resource through elevated proliferation rates; thus, we will refer to the phenotype that optimizes space consumption as **proliferators**. To understand eco-evolutionary dynamics resulting from selection for optimal phenotypic strategies toward creating (engineering) and consuming (proliferating) space, we consider an abstract on-lattice grid, where the domain is divided up into multiple subdomains by degradable ECM barriers.

Using a Cellular Automaton (CA) to implement our ABM, we examined the eco-evolutionary dynamics emerging under selection pressures that act to optimize mutually exclusive engineering and proliferating phenotypic strategies within domains with different spatial organization (C. Gatenbee et al., 2019; Margolus, n.d.; Poleszczuk and Enderling, 2014). We found that gaining access to new space through non-cell autonomous effects of engineers, can dramatically increase ITH and lead to patterns which, in the context of ecology, can be described as ecological succession. Surprisingly, we found that this selection-driven dynamic could lead, under sampling resolutions commonly used in cancer genomics, to subclonal diversification patterns that could be misinterpreted as evidence for neutral evolution.

## Results

### Model Description

We initiate simulations with 25 cells with the basic phenotype seeded in a subdomain, surrounded by ECM barriers. The subdomain itself is located in the center of the large domain, composed of the grid of subdomains. Cell growth is constrained to the subdomain, unless ECM separating the population from the neighboring subdomain is degraded. ECM barriers can be degraded through action of engineer cells if a sufficiently high concentration of ECM degrading enzymes is produced – which is a function of both the number of engineers as well as their efficiency (**Figure 1a**). When a cell divides, it can acquire a *driver* mutation that permanently increases either expression of ECM degrading enzymes (for the engineer phenotype), or proliferation rate (for proliferator phenotype). After each division, a check is made to remove any ECM which has met the removal criteria. A cell can die through apoptosis (with a rate of 15% each time step), freeing up space that can be explored by its neighbors. There is phenotype plasticity in the model, so engineers can become proliferators (at a rate of 0.2%) and vice-versa. Whereas these engineering and proliferative phenotypes are mutually exclusive, the mutational history of each cell enables them to start from the previous rates of engineering and proliferation if they switch back (**Figure 1b**). Simulations are initialized within an 601×601 domain, separated into smaller sub-domains, ranging from 25×25 to 301×301 (shown in **Figure 1c**), and run for 2500 timesteps, which allows for simulations to run for sufficiently long time so that the relevant evolutionary dynamics can be examined.

**Figure 1:**
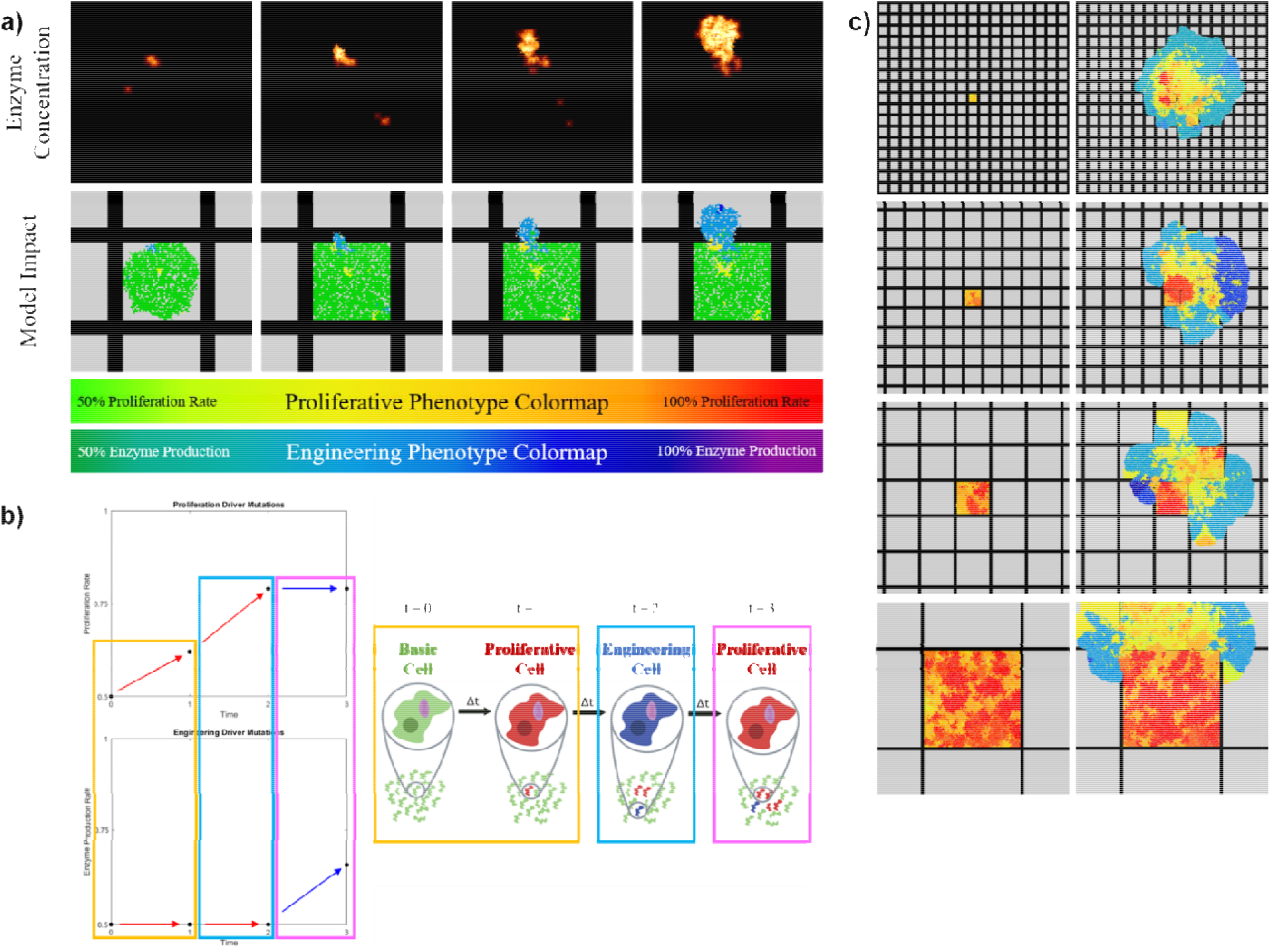
Model description. **a)** A depiction engineers degrading an ECM barrier in the model. For the ECM to be degraded, a sufficient concentration (~45% enzyme per lattice unit) of matrix-degrading enzyme is required near the ECM. The top images show the concentration of enzyme and the bottom images show the population of engineers producing them (with phenotypic color map below). **b)** A depiction of how driver mutations interact with phenotypic switching in the model. The top plot shows the proliferation driver mutation space, and the bottom plot shows the engineering driver mutation space. For all basic cells, their first driver mutation is randomly selected, which is the only time a mutation determines the cell’s phenotype (orange box). Afterwards, driver mutations will increase the cell’s phenotype at the time of the mutation. However, phenotypic switches occur at the end of a timeframe. The change boxed in blue shows the cell acquiring a proliferative driver (as it was the cell’s phenotype at the time of division), yet it stochastically switched phenotypes at the end of the timestep. Similarly, the cell acquires an engineering mutation in t = 3, and again stochastically switches phenotype at the end of the timestep (boxed in pink). The cell retains a memory of its mutational history, so the fitness does not reset upon switching phenotypes, yet only one phenotypic characteristic can be expressed at a time. **c)** Images taken from simulations of the four subdomain sizes in the model, showing the 2 types of characterized simulations. All parameters are the same except subdomain size. The left column shows simulations, where engineering failed, and the right column shows simulations where engineering succeeded.

### Impact of environmental engineering on phenotypic and mutational diversity

Regardless of subdomain size, we observed two scenarios characterized by engineering success. In some simulations, the engineering phenotype was driven to extinction, which left the population confined within the initial subdomain (failed engineering). In others, the population was capable of accessing the majority or all of the subdomains of the larger domain (**Figure 1c**). As a control, we examined the impact of domain size on phenotypic and clonal heterogeneity in the complete absence of engineering phenotype (**Figure S1a**). In the absence of successful engineering, phenotypic diversity peaked early, and then decreased as more effective proliferators were selected. Increase in the subdomain size lead to an increase in the peak phenotypic diversity. In the presence of successful engineering, phenotypic heterogeneity peaked at later time points, but at significantly higher levels. Similar to the outcome in the absence of engineering, the phenotypic diversity decreased after reaching its peak, indicating convergent evolution towards highest proliferation rates (**Figures 2a, S1a**). Clonal heterogeneity increased directly with subdomain size, reflecting an increase in population size (**Figures 2b, S1b**). After peaking, upon population reaching carrying capacity within spatial constrains, clonal heterogeneity remained high, with the effect more easily observed in larger subdomains. Successful engineering led to a delay in reaching maximal clonal heterogeneity but enabled populations to reach much higher levels of clonal heterogeneity. Notably, engineering limited reduction in clonal ITH, observed shortly after the peak. Therefore, environmental engineering resulting in production of public goods (space) enabled tumor populations, not only to gain access to space and achieve larger population size, but also increased levels of phenotypic and clonal heterogeneity.

**Figure 2:**
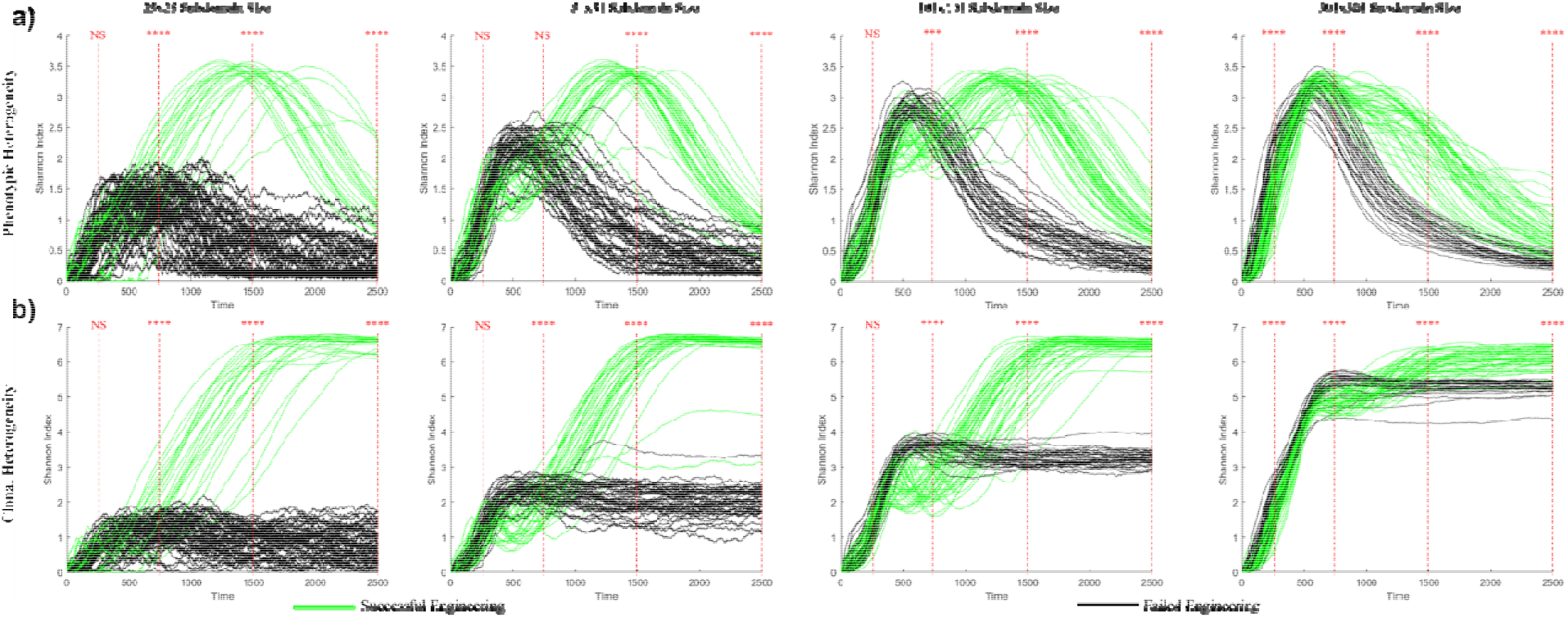
Impact of successful engineering on intra-tumor heterogeneity. Temporal changes in genetic **(a)** and phenotypic **(b)** heterogeneity are captured by Shannon Index. Green and black lines represent successful and failed engineering respectively. Each plot depicts the outcomes of 80 simulations. NS, *, **, *** and **** represent p values of Kolmogorov-Smirnov tests at the indicated time points: NS indicates p > 0.05, * indicates p<0.05, ** indicates p<0.01, *** indicates p<0.001, and **** indicates p<0.0001.

Successful engineering was associated with noticeable patterns of phenotypic succession. After breaking the initial ECM barrier, engineers exploited the newly accessed space, growing outwards the next barriers. As engineers continued to degrade ECM and grow outward, proliferative cells began to outcompete vestigial engineers near the core of the tumor (**Figure 3a, Supplemental Video 1**). Once all of the ECM barriers in the domain were removed, engineers were succeeded by proliferators, both from the expansion of pre-existing proliferative sub-clones and mutational conversion of engineers to proliferators (**Figures 3b, 3c**). Notably, this phenotypic succession was not accompanied by clonal one, as while phenotypic strategy converged toward proliferative phenotype, clonal heterogeneity was maintained at near peak levels. This result implies high levels of convergent evolution, where identical phenotypic solutions are reached by different subclonal lineages.

**Figure 3:**
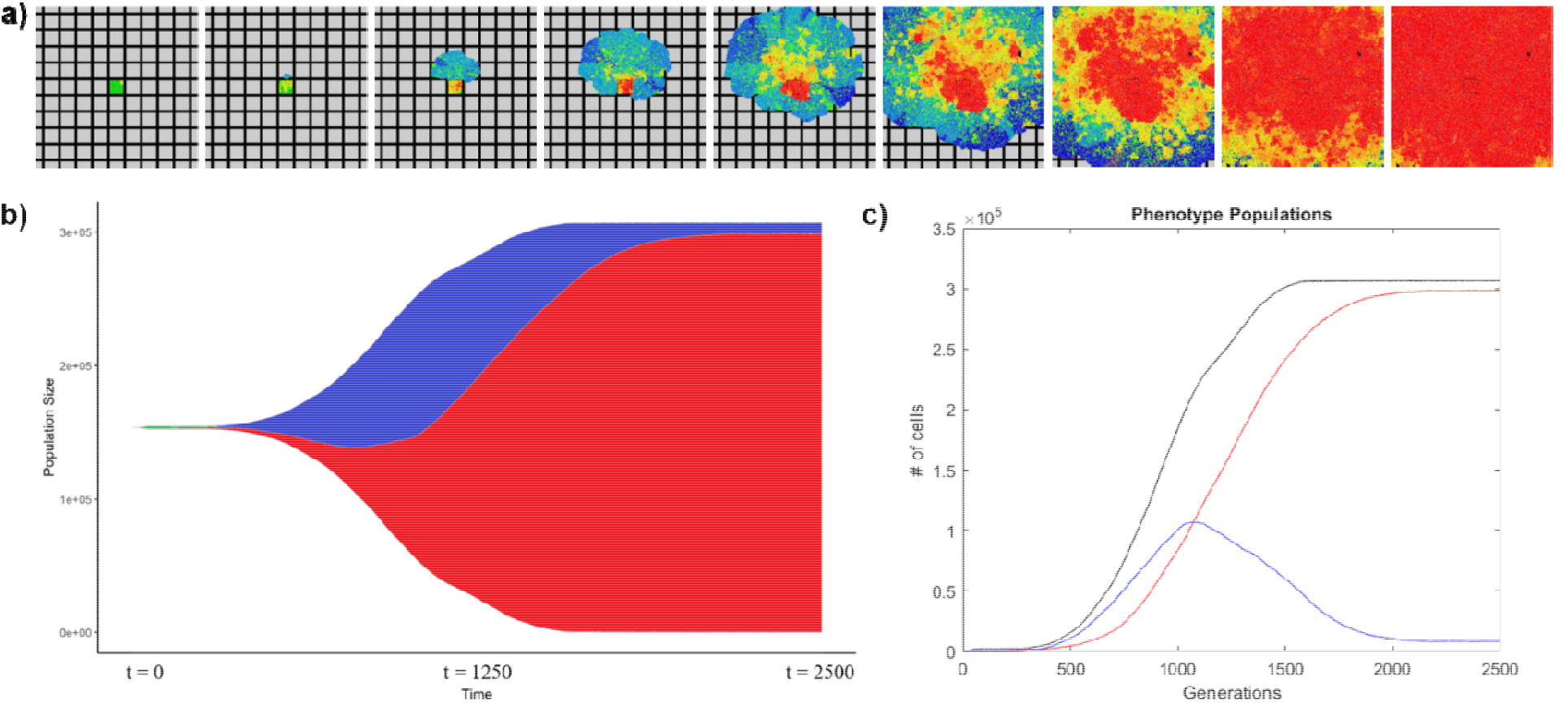
Patterns of ecological succession in the model. **a)** Images from the visualization of a simulation outcome showing engineers gaining access to new space and being succeeded by proliferative cells over time. **b)** Frequencies of basic, proliferative, and engineering cells over time. **c)** Populations of proliferative cells, engineering cells, and total tumor over time. Engineers flourish when they break out initially, and over time, proliferative cells can grow into the space created by engineers.

### Selection-driven dynamics can lead to clonal architectures consistent with neutral evolution

Since we recorded mutational history for each of the cell within simulation, this data enabled reconstruction of *true* (within simulations) clonal architecture, in contrast to inferred clonal architecture obtained from analyses of bulk genome sequencing. We visualized clonal architecture using Muller plots (C. D. Gatenbee et al., 2019). In this visualization, subclones with sub-threshold frequency (the majority of subclones in our simulations) are *invisible,* instead being grouped with the parental (sub)clone. We chose to visualize changes in clonal architecture using 10% clonal resolution threshold (**Figure 4a**), corresponding to a common 20x genome sequencing depth (assuming near diploid genome), and 1% clonal resolution threshold, reflecting higher resolution analyses (**Figure 4b**), and 0.1% threshold (**Figure S2**) – which provides a more accurate representation of the *true* clonal architecture.

**Figure 4:**
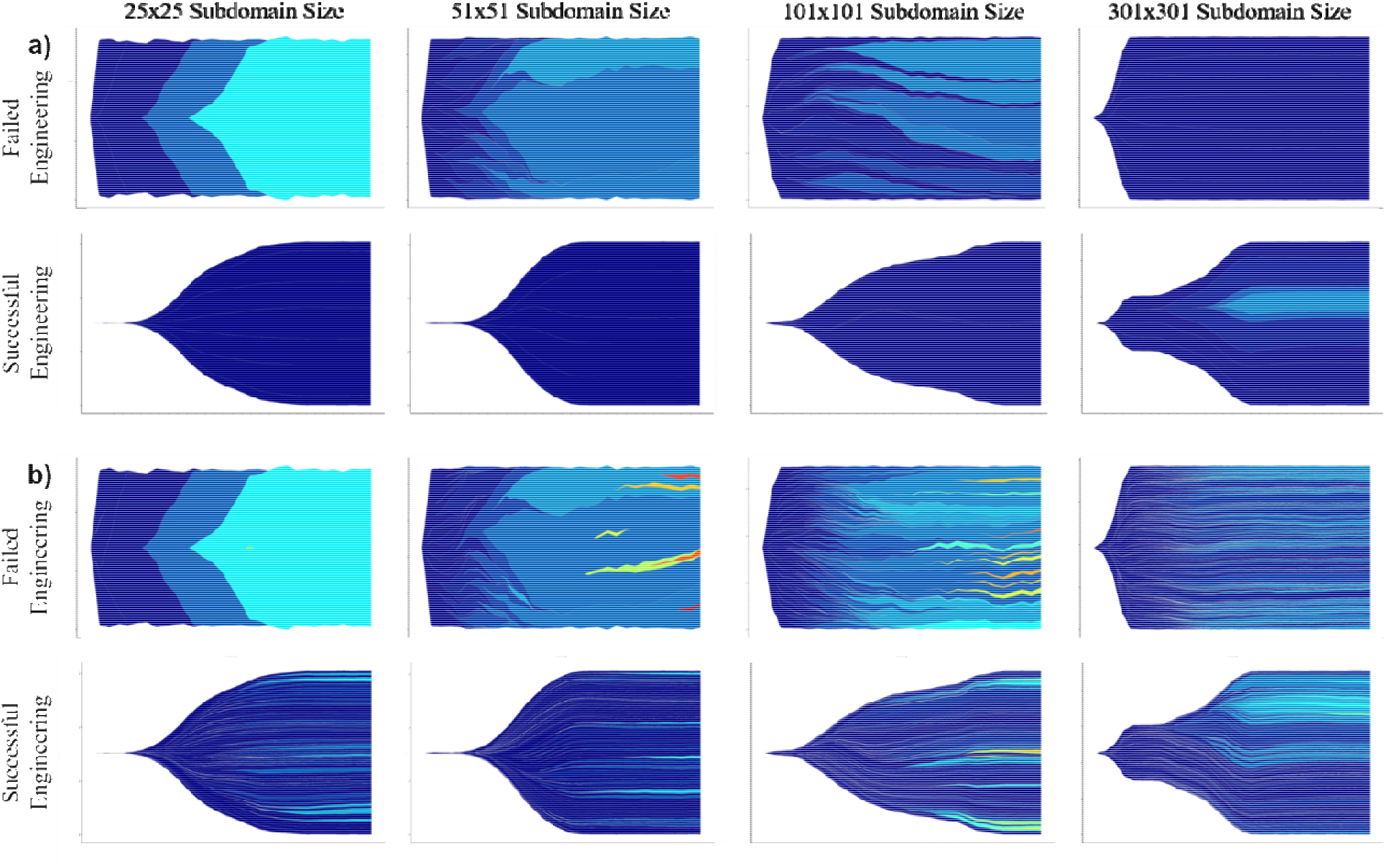
Muller plot visualization of the spatiotemporal clonal dynamics. Two representative simulations for each subdomain size are shown, one where engineering failed, and one where engineering succeeded. The x-axis on each plot is time, and the y-axis is population size. Colors indicate time of clone’s emergence, with earlier clones presented by dark blue, and later clones presented by warmer colors. Only clones with above-threshold frequency are plotted. **a)** Visualization with a frequency threshold of 10%. **b)** Visualization with a frequency threshold of 1%. Visualization with a frequency threshold of 0.1%is shown in **Figure S2**. Plots at different resolution levels represent same individual simulations.

In the absence of successful engineering, in the smallest subdomain size we observed patterns of clear clonal succession (**Figure 4**). Increase in the subdomain size resulted in tumors with more complex clonal architecture, but still with clear evidence of clonal expansions. Successful engineering dramatically increased clonal heterogeneity, leading to lower rates of expansion of individual sub-clones, indicative of increased clonal interference. Noticeably, increase of the analysis resolution revealed more complex clonal patterns (**Figure 4, Figure S2**).

Presence/absence of strong clonal expansions within the Muller’s plots visualization can be intuitively interpreted as evidence of selection-driven/neutral evolution. However, as such an interpretation might be misleading, we asked whether the apparent differences in patterns of clonal expansions observed at different resolution levels, would be interpreted differently when analyzed using quantitative metrics of neutrality. To this end, we examined the relation between the cumulative number of mutations as a function of the inverse frequency of the mutations. This approach has been used to discriminate between neutral (no fitness differences between subclonal lineages) and selection-driven evolution in inferences of evolution modes from bulk sequencing data, where a linear fit between the inverse of the variant allele frequencies (VAFs) of the detected mutations with the cumulative mutation function is interpreted as evidence of evolution neutrality (Williams et al., 2018, 2016). Since we had VAF data recorded from our simulations, we analyzed the goodness of linearity fit of our simulation data at different subdomain sizes and resolution levels of 1%, 0.1% and 0.01%. As a reference point, we generated VAF data from simulations that followed neutral dynamics (no changes in proliferation probability), under a scenario of no ECM barriers (lack of subdomains). Strikingly, we found a better linear fit of data from our selection-driven simulations, compared to the neutral control, at both 1% and 0.1% resolution, with the trend reversed at 0.01% (**Figure 5a**). The impact of resolution level on goodness of fit was not limited to our simulations, as we have also observed marked differences in the goodness of linearity fit within the artificial dataset provided with neutralitytestr (Williams et al., 2018, 2016) (**Figure S3**). Interestingly, comparison of the goodness of linearity fit between simulations with successful and failed engineering revealed significantly higher fit in simulations with successful engineering across all of the resolution levels, with the differences being more pronounced at lower subdomain size (**Figure 5b**). These results warrant caution in inferring mode of evolution from genomic sequencing data.

**Figure 5:**
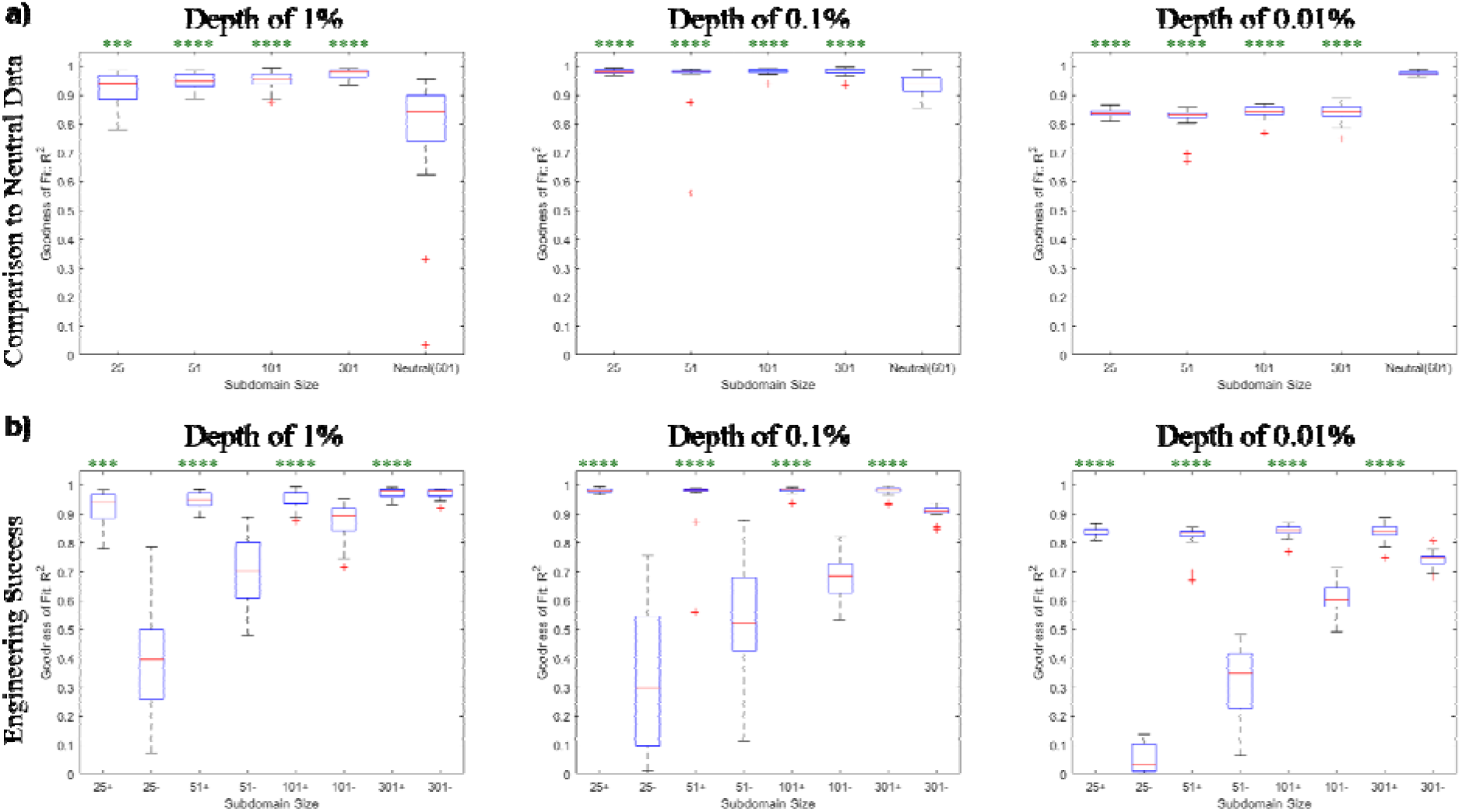
Consistency of clonal architecture with metrics of evolution neutrality. Goodness of fit to a linear regression between the cumulative mutation function and the inverse VAFs (M(f) 1/f) is used as a metrics of evolution neutrality. **a)** Goodness of feet for the in-silico growth simulations from Figure 2 with the indicated domain size and simulations with neutral growth dynamics. Statistical comparisons are made between simulations with successful engineering and the neutral simulations. **b)** Impact of engineering success on the goodness of fit. Statistical comparisons are made between the success and failure (indicated by + and – respectively) of each subdomain size. NS, *, **, *** and **** represent p values of Kolmogorov-Smirnov between two distributions: NS indicates p > 0.05, * indicates p<0.05, ** indicates p<0.01, *** indicates p<0.001, and **** indicates p<0.0001.

## Discussion

Our study examined the eco-evolutionary dynamics of tumor cell populations using an agent-based model where cells evolve under explicit pre-defined rules, specifying the impact of mutations and epigenetic switches on phenotypes that impact Darwinian competition for a limited ecological resource – space. In addition to the commonly used consideration of evolutionary fitness as a cell-intrinsic proliferation probability, our simulations included a second evolutionary strategy, which specified the ability of cells to unlock access to the *public good* of new space via non-cell autonomous action of environmental engineering, mediated by the action of secreted ECM remodeling enzymes. We found that non-cell autonomous production of common good had a profound impact on both phenotypic and clonal evolution, leading to higher heterogeneity and the emergence of patterns of phenotypic succession, while at the same time producing clonal architecture that might be consistent with a hypothesis of evolution neutrality.

Consistent with published reports of convergent evolution in cancers, where mutations with similar functional consequences are observed in distinct subclonal sub-populations (Ardaševa et al., 2019), we observed phenotypic convergence in our simulations, as different lineages within computational tumor were capable of finding optimal phenotypic solutions (Gerlinger et al., 2012). Whereas consideration of non-cell autonomous environmental engineering increased phenotypic diversity, it did not qualitatively impact convergence.

The addition of public good-enabled populations of tumor cells led to higher levels of clonal heterogeneity. To a large extent, this effect likely reflected the ability of these tumors to reach a much higher population size, as it was most pronounced within smaller subdomain size. However, public good engineering action also enabled a more effective maintenance of clonal ITH (notice the lack of a dip after reaching the peak in green versus black lines in **Figure 2a**) – consistent with the report of maintenance of clonal heterogeneity in a mouse model of non-cell autonomous driver of tumor growth (Marusyk et al., 2014).

The inclusion of producers of public good leads to more complex ecological dynamics, involving local and global phenotypic successions. The ability to degrade ECM barriers can, in some cases, lead to strong selection for engineering phenotypes, as they were capable of getting priority access to the newly available space – making the engineering strategy more effective at the growing edge of a tumor. However, as proliferative phenotypes were more successful in utilizing space, they eventually outcompeted engineers, leading to robustly reproducible patterns of phenotypic succession, which parallels ecological succession.

Even though the dynamic behavior in our simulations was designed to be governed by strong selection for competing phenotypic strategies, it produced patterns of clonal architecture that could be interpreted as indicative of neutral evolution patterns within current approaches towards inferences of evolution mode from genomic sequencing data. Most likely, this result reflects a lack of mutational record of history of phenotypic switching. Surprisingly, inferences of neutrality based on goodness of fit between the inverse of VAF and cumulative numbers of mutations showed higher consistency of the outcomes of our selection-driven simulations compared to simulations with explicitly neutral dynamics. Whereas increased resolution of mutation sampling led to superior fit in neutral simulations, the effect was only observed at resolution levels that exceed those employed in typical mutational analyses studies, with the exception of analyses of single crypts in colorectal cancers. Whereas our results do not disprove recent reports of neutral evolution in many cancers (Williams et al., 2018, 2016), they do warrant a more cautious interpretation of evolutionary inferences based on mutational analyses.

## Limitations of the Study

Whereas mathematical and computational modeling enables quantitative analysis of evolutionary dynamics in space in time, modeling deals with abstraction and simplification of biological reality. Moreover, the use of computational modeling depends on the accuracy of model assumptions and parameters, which are specified explicitly. While tumor growth requires gaining access to new space, this access is mediated not only by the action of tumor cells, but also by complex networks of microenvironmental interactions involving non-tumor cells, such as fibroblasts, macrophages etc. (Janiszewska et al., 2019; Marusyk et al., 2014). A concrete understanding of the underlying biology at this level, permitting adequate capture in a mathematical model is still missing. Even if such knowledge were available, quantitatively specifying all of relevant interactions would not be feasible. While primary tumors might be simpler than natural ecosystems, their ecological complexity is certainly higher than what can be captured in our models (Brown, 2016). Further, 2-dimensional lattices considered in our work do not faithfully recapitulate 3-dimensional tissue structures within primary or experimental tumors. On the other hand, computational modeling enables the examination of outcomes when the key ecological and evolutionary drivers of population dynamics, are considered – something which is not accessible not only to inferences from analyses of clinical samples, but also to experimental studies. Therefore, while quantitative results of our simulations might not be directly applicable to understanding of tumor biology, qualitative finding of the impact of ecological engineering on tumor heterogeneity and phenotypic succession is likely to be relevant to real tumors. Non-cell autonomous actions, including public goods, appear to be common in tumor ecosystems, including access to vasculature nutrients, immune invasion etc. (Tabassum and Polyak, 2015). Whereas our simulations only considered non-cell autonomous effects in terms of accessing spatial resources, results from our simulations are likely to be generalizable to other scenarios involving other public goods. We posit that consideration of ecological aspects of cancer evolution, the presence of non-cell autonomous effects, as well as integration of mathematical modeling with experimental studies and analyses of clinical samples will be essential to better understand clonal evolution –which is a prerequisite to the development of successful strategies to prevent and eradicate advanced cancers.

## Resource Availability

### Lead Contact

Further information and requests for resources and data should be directed to and will be fulfilled by the Lead Contact, Jack Edwards.

### Materials Availability

This study did not utilize any physical materials nor does it contain any biological data.

### Data and Code Availability

The model is publicly available on GitHub at CE_ABM: https://github/jackedwards1/CE_ABM

The data used to generate the figures is publicly available on Mendeley at http://dx.doi.org/10.17632/5wpcrnkvc7.1

## Acknowledgments

We would like to thank the Summer Program for the Advancement of Research Knowledge (SPARK) at the H. Lee Moffitt Cancer Center & Research Institute for sponsoring JE and the Integrated Mathematical Oncology Department at Moffitt for hosting this research project. We would also like to thank Dr. Joel Brown and Ryan Schenck for discussions and ideas. JE and DB were partially supported by an NCI IMAG-MSM grant (U01CA202958). DB was also funded by U01 CA244101. AM and DB were partially supported by the State of Florida award 20B06 (30-20450-99-01).

## Author Contributions

JE performed code development, data processing and statistical analyses. JE, AM and DB designed the study and wrote the manuscript. DB provided funding and overall supervision for the work.

## References

Altrock, P.M., Liu, L.L., Michor, F., 2015. The mathematics of cancer: integrating quantitative models. Nat. Rev. Cancer 15, 730–745. https://doi.org/10.1038/nrc4029

Ardaševa, A., Gatenby, R.A., Anderson, A.R.A., Byrne, H.M., Maini, P.K., Lorenzi, T., 2019. A mathematical dissection of the adaptation of cell populations to fluctuating oxygen levels (preprint). Cancer Biology. https://doi.org/10.1101/827980

Axelrod, R., Axelrod, D.E., Pienta, K.J., 2006. Evolution of cooperation among tumor cells. Proc. Natl. Acad. Sci. 103, 13474–13479. https://doi.org/10.1073/pnas.0606053103

Basanta, D., Anderson, A.R.A., 2013. Exploiting ecological principles to better understand cancer progression and treatment. Interface Focus 3, 20130020. https://doi.org/10.1098/rsfs.2013.0020

Bissell, M.J., Radisky, D., 2001. Putting tumours in context. Nat. Rev. Cancer 1, 46–54. https://doi.org/10.1038/35094059

Bissell, M.J., Radisky, D.C., Rizki, A., Weaver, V.M., Petersen, O.W., 2002. The organizing principle: microenvironmental influences in the normal and malignant breast. Differentiation 70, 537–546. https://doi.org/10.1046/j.1432-0436.2002.700907.x

Brown, J.S., 2016. Why Darwin would have loved evolutionary game theory. Proc. R. Soc. B Biol. Sci. 283, 20160847. https://doi.org/10.1098/rspb.2016.0847

Cawston, T.E., Wilson, A.J., 2006. Understanding the role of tissue degrading enzymes and their inhibitors in development and disease. Best Pract. Res. Clin. Rheumatol. 20, 983–1002. https://doi.org/10.1016/j.berh.2006.06.007

Chkhaidze, K., Heide, T., Werner, B., Williams, M.J., Huang, W., Caravagna, G., Graham, T.A., Sottoriva, A., 2019. Spatially constrained tumour growth affects the patterns of clonal selection and neutral drift in cancer genomic data (preprint). Bioinformatics. https://doi.org/10.1101/544536

Gatenbee, C., West, J., Baker, A.M., Guljar, N., Jones, L., Graham, T.A., Robertson-Tessi, M., Anderson, A.R.A., 2019. Macrophage-mediated immunoediting drives ductal carcinoma evolution: Space is the game changer (preprint). Cancer Biology. https://doi.org/10.1101/594598

Gatenbee, C.D., Schenck, R.O., Bravo, R.R., Anderson, A.R.A., 2019. EvoFreq: visualization of the Evolutionary Frequencies of sequence and model data. BMC Bioinformatics 20, 710. https://doi.org/10.1186/s12859-019-3173-y

Gatenby, R.A., Gillies, R.J., 2008. A microenvironmental model of carcinogenesis. Nat. Rev. Cancer 8, 56–61. https://doi.org/10.1038/nrc2255

Gerlinger, M., Rowan, A.J., Horswell, S., Larkin, J., Endesfelder, D., Gronroos, E., Martinez, P., Matthews, N., Stewart, A., Tarpey, P., Varela, I., Phillimore, B., Begum, S., McDonald, N.Q., Butler, A., Jones, D., Raine, K., Latimer, C., Santos, C.R., Nohadani, M., Eklund, A.C., Spencer-Dene, B., Clark, G., Pickering, L., Stamp, G., Gore, M., Szallasi, Z., Downward, J., Futreal, P.A., Swanton, C., 2012. Intratumor Heterogeneity and Branched Evolution Revealed by Multiregion Sequencing. N. Engl. J. Med. 366, 883–892. https://doi.org/10.1056/NEJMoa1113205

Greaves, M., Maley, C.C., 2012. Clonal evolution in cancer. Nature 481, 306–313. https://doi.org/10.1038/nature10762

Hanahan, D., Weinberg, R.A., 2011. Hallmarks of Cancer: The Next Generation. Cell 144, 646–674. https://doi.org/10.1016/j.cell.2011.02.013

Hanahan, D., Weinberg, R.A., 2000. The Hallmarks of Cancer. Cell 100, 57–70. https://doi.org/10.1016/S0092-8674(00)81683-9

Janiszewska, M., Tabassum, D.P., Castaño, Z., Cristea, S., Yamamoto, K.N., Kingston, N.L., Murphy, K.C., Shu, S., Harper, N.W., Del Alcazar, C.G., Alečković, M., Ekram, M.B., Cohen, O., Kwak, M., Qin, Y., Laszewski, T., Luoma, A., Marusyk, A., Wucherpfennig, K.W., Wagle, N., Fan, R., Michor, F., McAllister, S.S., Polyak, K., 2019. Subclonal cooperation drives metastasis by modulating local and systemic immune microenvironments. Nat. Cell Biol. 21, 879–888. https://doi.org/10.1038/s41556-019-0346-x

Kagel, J.H., Roth, A.E., 1995. The Handbook of Experimental Economics. Princeton University Press.

Korolev, K.S., Xavier, J.B., Gore, J., 2014. Turning ecology and evolution against cancer. Nat. Rev. Cancer 14, 371–380. https://doi.org/10.1038/nrc3712

Lloyd, M.C., Cunningham, J.J., Bui, M.M., Gillies, R.J., Brown, J.S., Gatenby, R.A., 2016. Darwinian Dynamics of Intratumoral Heterogeneity: Not Solely Random Mutations but Also Variable Environmental Selection Forces. Cancer Res. 76, 3136–3144. https://doi.org/10.1158/0008-5472.CAN-15-2962

Margolus, N., n.d. Cellular Automata Machines 27.

Marusyk, A., Tabassum, D.P., Altrock, P.M., Almendro, V., Michor, F., Polyak, K., 2014. Non-cell-autonomous driving of tumour growth supports sub-clonal heterogeneity. Nature 514, 54–58. https://doi.org/10.1038/nature13556

Myers, K.V., Pienta, K.J., Amend, S.R., 2020. Cancer Cells and M2 Macrophages: Cooperative Invasive Ecosystem Engineers. Cancer Control 27, 107327482091105. https://doi.org/10.1177/1073274820911058

Noble, R., Burri, D., Kather, J.N., Beerenwinkel, N., 2019. Spatial structure governs the mode of tumour evolution (preprint). Cancer Biology. https://doi.org/10.1101/586735

Nowell, P., 1976. The clonal evolution of tumor cell populations. Science 194, 23–28. https://doi.org/10.1126/science.959840

Poleszczuk, J., Enderling, H., 2014. A High-Performance Cellular Automaton Model of Tumor Growth with Dynamically Growing Domains. Appl. Math. 05, 144–152. https://doi.org/10.4236/am.2014.51017

Sottoriva, A., Kang, H., Ma, Z., Graham, T.A., Salomon, M.P., Zhao, J., Marjoram, P., Siegmund, K., Press, M.F., Shibata, D., Curtis, C., 2015. A Big Bang model of human colorectal tumor growth. Nat. Genet. 47, 209–216. https://doi.org/10.1038/ng.3214

Swanton, C., 2012. Intratumor Heterogeneity: Evolution through Space and Time. Cancer Res. 72, 4875–4882. https://doi.org/10.1158/0008-5472.CAN-12-2217

Tabassum, D.P., Polyak, K., 2015. Tumorigenesis: it takes a village. Nat. Rev. Cancer 15, 473–483. https://doi.org/10.1038/nrc3971

West, J., Schenck, R.O., Gatenbee, C., Robertson-Tessi, M., Anderson, A.R.A., 2019. Tissue structure accelerates evolution: premalignant sweeps precede neutral expansion (preprint). Cancer Biology. https://doi.org/10.1101/542019

Williams, M.J., Werner, B., Barnes, C.P., Graham, T.A., Sottoriva, A., 2016. Identification of neutral tumor evolution across cancer types. Nat. Genet. 48, 238–244. https://doi.org/10.1038/ng.3489

Williams, M.J., Werner, B., Heide, T., Curtis, C., Barnes, C.P., Sottoriva, A., Graham, T.A., 2018. Quantification of subclonal selection in cancer from bulk sequencing data. Nat. Genet. 50, 895–903. https://doi.org/10.1038/s41588-018-0128-6

