## Supplemental Information for "Selection-driven tumor evolution involving non-cell growth promotion leads to patterns of clonal expansion consistent with neutrality interpretation"

**Supplemental Figures**

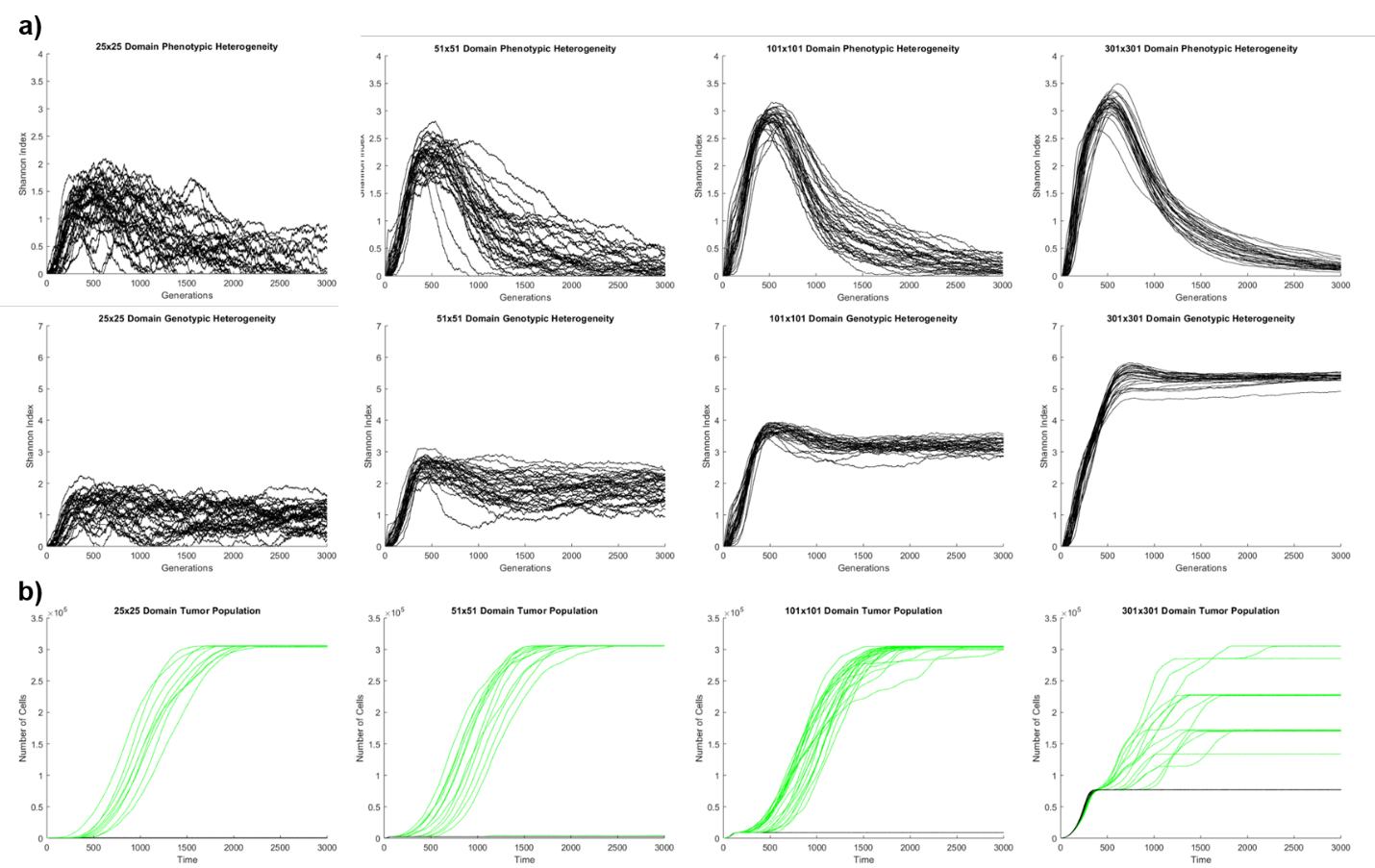

**Supplemental Figure 1**: **Control data and population data. a)** Phenotypic heterogeneity (top) and clonal heterogeneity (bottom) for simulations over time when the engineering phenotype was disabled. There are 30 simulations per plot, each run for 3000 timesteps. **b)** Total tumor population over time in simulations with the same parameters as Figure 2. Black lines represent failed engineering, green lines represent successful engineering. The failed engineering tumor carrying capacity is dependent on what the subdomain size is.

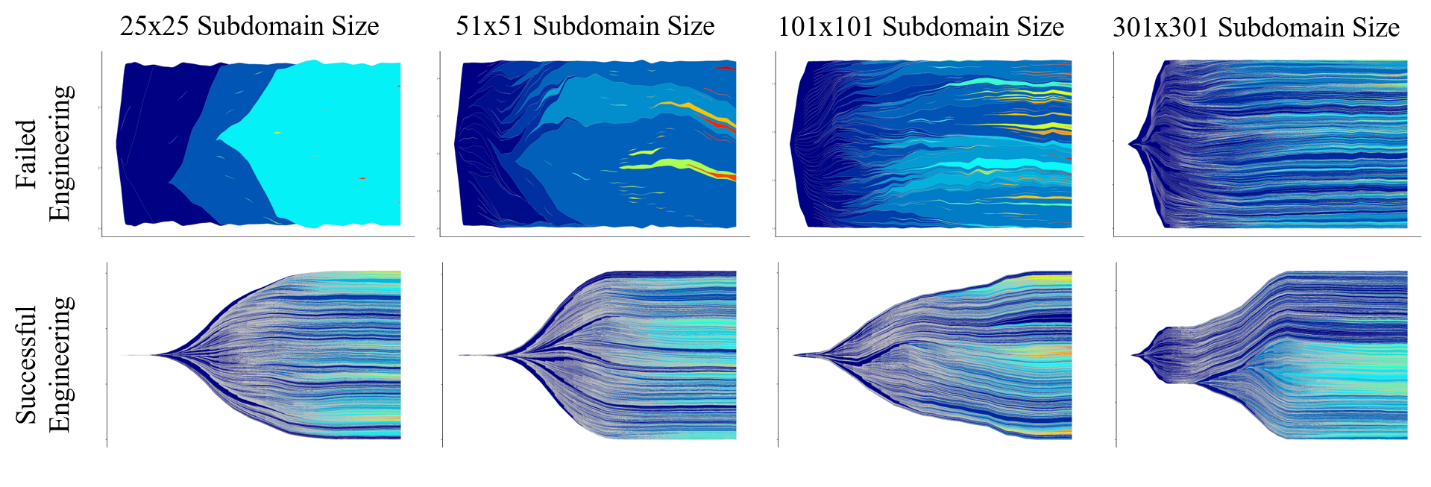

**Supplemental Figure 2**: **Muller plot visualization of the spatiotemporal clonal dynamics at 0.1% resolution.** Two representative simulations for each subdomain size are shown (same simulations as in Figure 4), one where engineering failed, and one where engineering succeeded. The x-axis on each plot is time, and the y-axis is population size. Colors indicate time of clone’s emergence, with earlier clones presented by dark blue, and later clones presented by warmer colors. Only clones with above-threshold frequency are plotted.

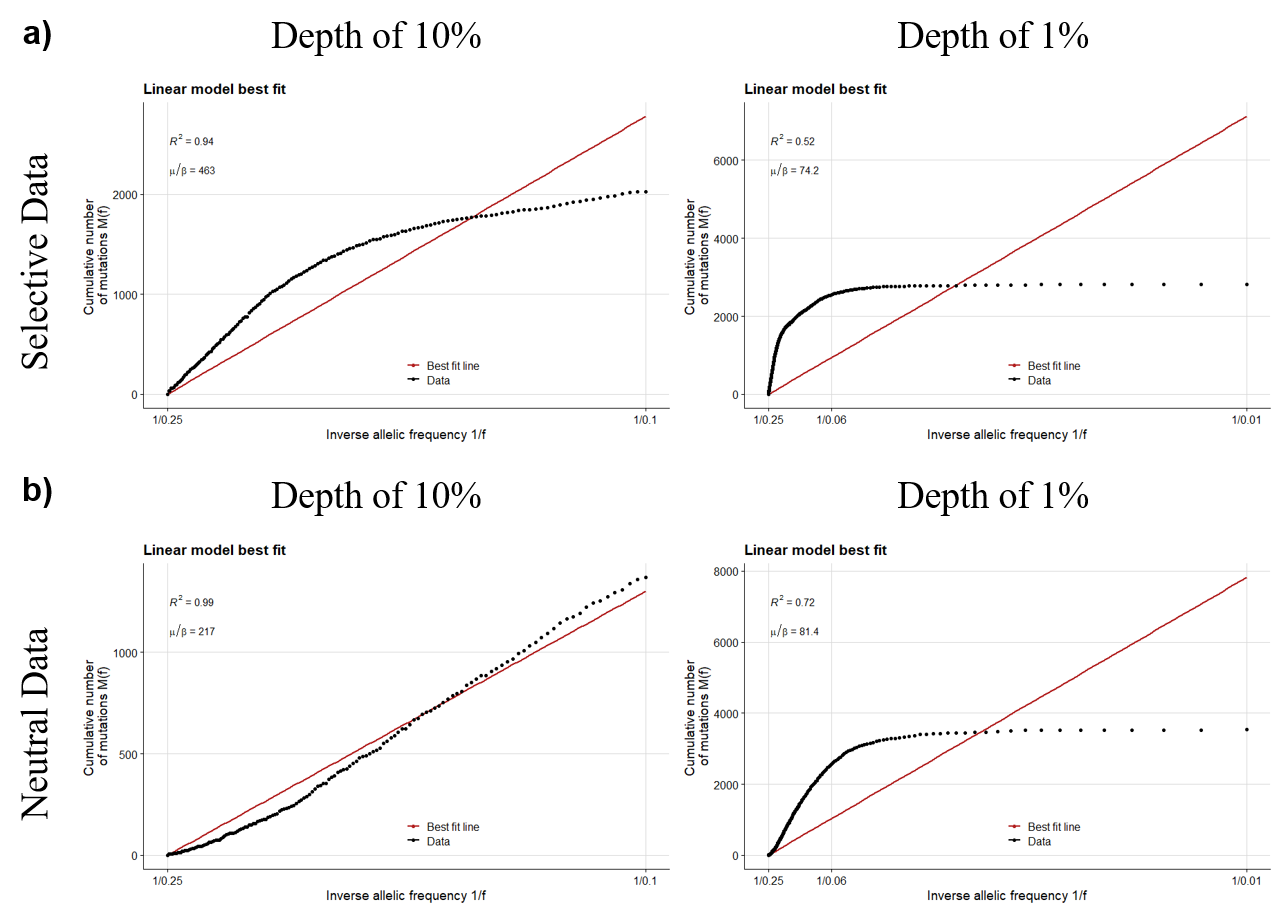

**Supplemental Figure 3:** **Examining the effect of frequency on the inverse allelic frequency metric for neutrality.** Artificial data that comes with the R package neutralitytestr (Williams et al., 2018, 2016). The plots show the cumulative number of mutations as a function of the inverse allelic frequency, and the goodness of fit to a linear distribution, as a metrics for a neutral mode of evolution. **a)** The selective data plotted from frequencies of 25% to 10%. **b)** The neutral data plotted from frequencies of 25% to 1%.

**Transparent Methods**

The model uses a 2D on-lattice agent-based model, where each grid point is either unoccupied, blocked by ECM, or is occupied by a living cell. There are three cell phenotypes: basic, proliferative, and engineering. Cells with the basic phenotype (basic) are capable of proliferating while ignoring homeostatic signaling constraints, as long as space is available, but are incapable of accessing new space through degradation of matrix. Cells with the engineering phenotype (engineers) proliferate at the same rates as basic cells but can provide access to new space by degrading ECM barriers through secretion of matrix degrading enzymes. No single engineer can produce enough of the enzyme on its own to degrade the ECM; a substantial population of engineers local to the ECM must secrete enzymes at the same time to access new space. Cells with a proliferative phenotype (proliferators) can divide at higher rates than basic and engineer cells, thus they are more effective at claiming available space, but they cannot gain access to new space on their own.

The CA uses a Moore neighborhood to define the cells in direct contact with each other (Margolus, n.d.). Cells are iterated over stochastically as to avoid artifacts or giving preference to cells in specific spatial locations. Each cell determines if it has the ability (fitness) and resources (space) to divide during that timestep. Cells have a potential proliferation rate which defines its ability to proliferate if space is available. For instance, a cell with a potential proliferation rate of 75% has a 75% probability to divide at each time step, provided there is space. Potential proliferation starts at 50% within the basic phenotype and can be increased by mutations that optimize proliferative phenotype – up to 100% (dividing at each time step given space availability). At any given timestep, a cell has a 15% probability of dying, which leaves an empty space in the next timestep, enabling cell turnover within the core of the tumor.

At each cell division, one of the following two types of *driver* mutations might occur at a total probability of 0.07%. First, a proliferation-driving mutation (0.07% probability), can increase cell proliferation with a randomly chosen increment of 0.01% between 10 and 20%. Second, engineering-driving mutation, occurring at 0.07% probability, bestows a cell with an engineering phenotype, enabling the mutated cell to produce a matrix-degrading enzyme at 50% probability per time step. The enzyme-production ability of engineering phenotypes can be further increased by additional engineering mutations, through increments of 0.01% in the 10 - 20% range. In addition to driver mutations, *passenger* mutations, occurring with 0.5% per time step probability, could be either entirely neutral (no impact on cell fitness), or slightly deleterious by decreasing probability of proliferation in the range 0 to 0.5%, with 0.01% increment. If no mutations occur, a cell retains its current phenotype, which is inherited upon cell division.

When a basic cell first acquires a driver mutation, it will randomly become either a proliferator cell or an engineer. Further driver mutations will be chosen dependent on the cell’s current phenotype; proliferative cells will always acquire proliferative drivers, and engineering cells will always acquire engineering drivers. Based on the principle of evolutionary tradeoff, we consider engineer (ECM-degrading) or proliferative phenotypes to be mutually exclusive (Townsend et al., 2009). However, a cell can switch its phenotype from engineer to proliferator and vice versa at a probability of 0.2% at the end of each timestep. This type of stochastic phenotypic switching among cancer cell has been described in the literature (Gupta et al., 2011; Zhou et al., 2014). When a phenotypic switch occurs, we assume that the cell loses the expression of improved abilities related to the pre-switch phenotype. If this cell or its clonal progeny undergo a reverse switch, it will recover these abilities at the levels that preceded the first switch. As an example, an engineer cell that undergoes phenotypic switch toward proliferator will stop degrading ECM and will proliferate at a rate that would correspond to be basal one if it never had a proliferative phenotype before. If it subsequently undergoes a mutation that transforms it into a cell with an engineer phenotype, the ability to degrade ECM will be restored to the level it had previously as an engineer.

In order to interrogate the accumulation of *genetic* ITH, we record mutational history for each cell. Each driver mutation or any successive four passenger mutations were used in the analyses as a mutational branching point, marking a new sub-clone. *Phenotypic* strategies and mutational history were tracked for all the lineages over time. To define phenotypic ITH, we binned cells based on their proliferation and enzyme-producing phenotypes, with increments of 1%. Thus, for example, two proliferator cells representing different genetic subclonal branches, with proliferation probabilities of 59.1% and 59.7% would be counted as belonging to the same *phenotypic* subpopulation. Based on this definition of phenotypic subpopulations, there were 103 possible distinct phenotypic groups: 51 proliferative, 51 engineering, and 1 basic. To visualize outcomes of the simulations, we map a color to a cell’s phenotype and fitness as described by the aforementioned rules. This map is visible in **Figure 1a**.

A concentration of ~45% enzyme per unit lattice is required local to a point of ECM to remove it in the model. The enzyme produced by engineers is considered to be a diffusible molecule that decays over time. In order for an ECM barrier to be degraded successfully, there needs to be at least 5 engineers producing enzyme immediately next to the barrier, and the minimum number of engineers increases with distance away from the ECM. The maximum diffusion range on the enzyme is a radius of 3 cells out from the grid point where the engineer sits, and the effectiveness follows a simple linear decay in effectiveness (as opposed to an inverse square law). Successful destruction of ECM unlocks the space within the empty subdomain. The newly accessed space resource can be exploited by all cell types.

The parameters chosen for these simulations were selected with the goal that they would lead to realistic looking simulations. A wide variety of parameters were tested, both during and after model development and yielded similar outcomes, convincing us that our results are not just an artifact of the chose parameters. We varied mutation rate [0.005% - 1%], impact of driver mutations [2.5% - 25%], death rate [5% - 50%], and enzyme strength [~20% - ~80%]. The parameters used in the simulations we show are optimal for ensuring all of the relevant eco-evolutionary dynamics play out within a reasonable timeframe, as well as consistently getting a roughly even split of engineering fails and successes in order to make comparisons between the two simulations. The specific parameters used in the reported simulations are: a total simulation length of 2500 timesteps, a mutation bias of 0 (engineering and proliferation mutations are equally likely), a passenger mutation rate of 0.5%, a driver mutation rate of 0.07%, a switching frequency of 0.2%, a driver impact range of 10.5% to 20%, a death rate of 15%, an enzyme radius of 2, and an integer enzyme threshold for degradation of 10 (a strength of 10 enzymes must be present local to a piece of ECM in order for it to be removed).

When computing heterogeneity for both the phenotypic groups and the clonal groups, the Shannon index was used (Shannon, n.d.). The model takes the proportion of each of the phenotypic populations, *p_i_*, and calculates the Shannon Index as follows:

$$H= -\sum_{i=1}^{R} p_{i}*ln(p_{i})$$

*H* was calculated for the phenotypic groups and the clones to measure the differences in clonal heterogeneity and the phenotypic diversity of the tumor. A Kolmogorov-Smirnov test was performed to examine the quantitative differences in the heterogeneity distributions between simulations characterized by successful and failed engineering. During the simulation, a list of all clones and their ancestry is recorded and output. From this master list, muller plots can be constructed and most non-spatial neutrality statistics can be computed. In the data processing, we used the master list for each simulation to create a long-format output of the clonal architecture, which is readable by EvoFreq, an R package designed for visualizing clonal dynamics (Gatenbee et al., 2019). We also used the master list to calculate the variant allelic frequencies (VAFs). For our neutrality metric, we compared the cumulative mutations as function of the inverse frequency to a linear model, which is theoretically a neutral mode of growth (Williams et al., 2018, 2016). It is shown that the cumulative amount of distributions, *M(f)*, is described by the following equation in a neutral model of evolution:

$$M\left( f \right)= \frac{\mu}{\beta}(\frac{1}{f}-\frac{1}{f_{max}})$$

where *f* is the variant allelic frequency of a given mutation, *f_max_* is the maximum frequency of a mutation, *µ* is the mutation rate, and *β* is the cell division rate in which both lineages survive (Williams et al., 2016). The parameters and derivation of this model are detailed explicitly by Williams et al. in 2016. We examined the goodness of fit of our data to a linear (neutral) model, and also compared our data to both an explicitly neutral simulation in our model as well as the artificial data provided in an R package, neutralitytestr, that is designed to test VAF data using the linear fit method described (Williams et al., 2018, 2016). Again, a Kolmogorov-Smirnov test was performed to quantify the differences between the distributions of our model data and explicitly neutral simulations.

**Supplemental References**

Gatenbee, C.D., Schenck, R.O., Bravo, R.R., Anderson, A.R.A., 2019. EvoFreq: visualization of the Evolutionary Frequencies of sequence and model data. BMC Bioinformatics 20, 710. https://doi.org/10.1186/s12859-019-3173-y

Gupta, P.B., Fillmore, C.M., Jiang, G., Shapira, S.D., Tao, K., Kuperwasser, C., Lander, E.S., 2011. Stochastic State Transitions Give Rise to Phenotypic Equilibrium in Populations of Cancer Cells. Cell 146, 633–644. https://doi.org/10.1016/j.cell.2011.07.026

Margolus, N., n.d. Cellular Automata Machines 27.

Shannon, C.E., n.d. A Mathematical Theory of Communication 55.

Townsend, C.R., Begon, M., Harper, J.L., 2009. Essentials of Ecology. [electronic resource]., 3rd ed. ed. John Wiley & Sons, Ltd.

Williams, M.J., Werner, B., Barnes, C.P., Graham, T.A., Sottoriva, A., 2016. Identification of neutral tumor evolution across cancer types. Nat. Genet. 48, 238–244. https://doi.org/10.1038/ng.3489

Williams, M.J., Werner, B., Heide, T., Curtis, C., Barnes, C.P., Sottoriva, A., Graham, T.A., 2018. Quantification of subclonal selection in cancer from bulk sequencing data. Nat. Genet. 50, 895–903. https://doi.org/10.1038/s41588-018-0128-6

Zhou, J.X., Pisco, A.O., Qian, H., Huang, S., 2014. Nonequilibrium Population Dynamics of Phenotype Conversion of Cancer Cells. PLoS ONE 9, e110714. https://doi.org/10.1371/journal.pone.0110714
