## Supplementary figures and images for "Selection-driven tumor evolution involving non-cell growth promotion leads to patterns of clonal expansion consistent with neutrality interpretation"

### Supplementary Video 1

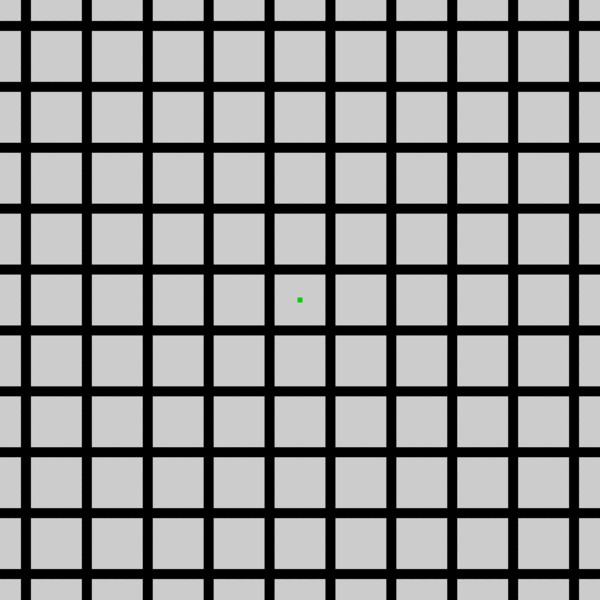
